# The Behavior of Molecular Measures of Natural Selection after a Change in Population Size

**DOI:** 10.1101/2020.12.18.423515

**Authors:** Rebekka Müller, Ingemar Kaj, Carina F. Mugal

**Affiliations:** Department of Mathematics, Uppsala University, 752 37 Uppsala, Sweden; Department of Ecology and Genetics, Uppsala University, 752 36 Uppsala, Sweden; Norbyvägen 18D, 752 36 Uppsala, Sweden

**Keywords:** theoretical population genetics, nearly neutral theory, natural selection, genetic drift, effective population size

## Abstract

A common model to describe natural selection at the molecular level is the nearly neutral theory, which emphasizes the importance of mutations with slightly deleterious fitness effects as they have a chance to get fixed due to genetic drift. Since genetic drift is stronger in smaller than in larger populations, a negative relationship between molecular measures of selection and population size is expected within the nearly neutral theory. Originally, this hypothesis was formulated under equilibrium conditions. A change in population size, however, pushes the selection-drift balance off equilibrium leading to alterations in the efficacy of selection. To investigate the nonequilibrium behavior, we relate measures of natural selection and genetic drift to each other, considering both, measures of micro- and macroevolution. Specifically, we use a Poisson random field framework to model *π_N_/π_S_* and *ω* as time-dependent measures of selection and assess genetic drift by an effective population size. This analysis reveals a clear deviation from the expected equilibrium selection-drift balance during nonequilibrium periods. Moreover, we find that microevolutionary measures quickly react to a change in population size and reflect a recent change well, at the same time as they quickly lose the knowledge about it. Macroevolutionary measures, on the other hand, react more slowly to a change in population size but instead capture the influence of ancient changes longer. We therefore conclude that it is important to be aware of the different behaviors of micro- and macroevo- lutionary measures when making inference in empirical studies, in particular when comparing results between studies.

## INTRODUCTION

Among the key driving factors of evolution are mutations, selection, and genetic drift. The analysis of the interplay between them therefore provides valuable understanding on a population’s ability to evolve and adapt. Population genetics theory predicts that the strength of genetic drift is weaker in larger populations than in smaller populations, due to the stochastic nature of reproduction (Wright 1931; Kimura 1964). This results in a positive correlation between the efficacy of selection and population size, and hence deleterious mutations are more likely to go extinct due to purifying selection if population size is large. This relationship can for example explain why asexual (Lynch *et al.* 1993; Lumley *et al.* 2015), small, or endangered populations (Lynch *et al.* 1995; Lachapelle *et al.* 2017) have a higher risk to go extinct or suffer from genetic diseases (Karlsson *et al.* 2014; Seppälä 2015).

The relationship between the efficacy of selection and population size also provides the foundation for the nearly neutral theory of molecular evolution (Ohta 1973, 1976, 1992). In this framework, typically the distribution of fitness effects (DFE) of new mutations is weighted towards purifying selection: most mutations are deleterious, of which a nonnegligible amount is slightly deleterious, and only a small proportion of mutations is advantageous. While strongly deleterious mutations rarely segregate in the population or get fixed, slightly deleterious mutants contribute significantly to segregating polymorphisms and can eventually get fixed due to genetic drift. The fact that only a small fraction of mutants is under positive selection makes the nearly neutral theory predict a negative correlation between molecular measures of natural selection and population size. The validity of this hypothesis can be tested by comparing measures of selection and genetic drift, and has been supported across a wide range of taxa (e.g. Hughes 2008; Elyashiv *et al.* 2010; Figuet *et al.* 2016; Chen *et al.* 2017; Bolívar *et al.* 2019).

The traditional approach to assessing genetic drift is to apply some concept of an effective population size, *N*_eff_. The concept of *N*_eff_ is used to compare a given (non-ideal) population with a simpler idealized reference model, such as the ideal Wright-Fisher model, with respect to a particular property. This leads to different definitions of *N_eff_,* e.g. inbreeding and variance (Wright 1931; Crow and Kimura 1970) or eigenvalue effective size (Ewens 1979). All approaches predict *N*_eff_ under different circumstances as for example certain spatial and temporal scales and demographic scenarios. Most often it is not evident whether underlying assumptions of the various models are met and how accurate resulting estimates of *N*_eff_ are when assumptions are violated. For this, the spatial and temporal scales of different estimates of *N*_eff_ have to be interpreted (Wang *et al.* 2016) to draw the right conclusions. These issues are discussed frequently in the literature (Ewens 1969; *Sjödin et al.* 2004; Waples 2005; Wang *et al.* 2016; Galtier and Rousselle 2020) confirming the need for more clarification in this field.

To detect evidence of selection in genome data, different approaches and methods have been developed. A common feature of them is to contrast neutral reference and test data. Under the assumption that synonymous mutations are neutral, the contrast between synonymous and nonsynony- mous mutations in protein-coding sequences provides a relevant approach. Measures of natural selection at the microevolutionary timescale, which are based on polymorphisms, are designed to identify selective events within a species and give a *snapshot* of the current state. A popular representative is the ratio of nonsynonymous and synonymous diversity, *π_N_/π_S_* (Nei and Li 1979). Other measures look at interspecific differences in form of fixations that arise through events which happened in a lineage since a speciation event. Those measures are *accumulative* and give insight into the macroevolution of a population or species. A measure that belongs to this group is the ratio of the nonsynonymous and synonymous fixation rate, ω, commonly assessed by the ratio of nonsynonymous and synonymous sequence divergence (Goldman and Yang 1994; Muse and Gaut 1994).

The nearly neutral theory predicts a negative correlation between *N*_eff_ and the measures *π_N_/π_S_* and *ω*: the smaller *N*_eff_ the larger the ratios, i.e. the less efficient selection. However, the prediction is based on the equilibrium assumption, where the effect of genetic drift on segregating sites balances the efficacy of selection in a stable manner implying a constant evolutionary rate. Yet there are many factors that can cause a nonequilibrium such as a change in population size, population structure or migration, a change in recombination rate, or environmental variation (Brandvain and Wright 2016). These events disturb the selection-drift balance.

Several empirical studies (Lu *et al.* 2006; Campos *et al.* 2014; Hollister *et al.* 2014; Renaut and Rieseberg 2015; Leroy *et al.* 2020; Galtier and Rous- selle 2020) provide evidence that the efficacy of natural selection is limited in nonequilibrium conditions. In a meta-analysis, Brandvain and Wright (2016) compare predictions of classical theory with results from a large number of data analyses, which stresses the need of care for nonequilibrium conditions when testing for differences in selection efficacy between species. To enable such care to be taken, theoretical models are a critical tool to provide conceptual understanding of the interplay between selection and genetic drift and the extent of nonequilibrium periods. Given the mathematical challenges in modeling natural selection in nonequilibrium scenarios, such studies are still rare. Gordo and Dionisio (2005), for example, develop a nonequilibrium model to estimate parameters of deleterious mutations, which can be applied to quantify the mutation rate to deleterious alleles. Alternatively, simulations can be used to provide observational insight into the nonequilibrium behavior of measures of evolution. Rousselle *et al.* (2018) investigate the rate of adaptive molecular evolution in fluctuating population size and find that this violation of the equilibrium assumption can lead to a severely biased inference based on current methodology.

In this study we investigate the effect of a change in population size on micro- and macroevolutionary measures of selection in an otherwise ideal population. While being aware of other factors that lead to a nonequilibrium, concentrating on this isolated aspect provides conceptual understanding that is straight forward to interpreted. For this purpose, we first introduce a mathematical model and state and derive relevant quantities needed to model measures of selection in nonequilibrium. Among these, an important statistic is the allele frequency spectrum (AFS) from which other measures such as nucleotide diversity can be derived. Several different approaches exist to model the AFS in nonequilibrium (Evans *et al.* 2007; Živković and Stephan 2011; Živković *et al.* 2015; Kaj and Mugal 2016). We here build on the Poisson random field framework approach as in Kaj and Mugal (2016). With the help of this framework, the above addressed measures, *π_N_/π_S_* and *ω*, are formulated as functions of time, (*π_N_/π_S_*)(*t*) and *ω*(*t*). Equipped with the analytical model, we discuss the following questions. First, how does a change in population size affect micro- and macroevolutionary measures of natural selection, (*π_N_/π_S_*)(*t*) and *ω*(*t*)? Second, does the prediction of the nearly neutral theory also hold in a nonequilibrium? To this end we investigate different choices of *N*_eff_ as measures of genetic drift. Finally, we discuss the relevance of micro- and macroevolutionary measures in the study of natural selection.

## MODELS AND METHODS

### Model formulation

Let *N* represent the population size of a haploid population—which under the assumption of additive fitness effects in a diploid organism, is equivalent to a diploid population of size *N*/2. We consider the evolution of a population that undergoes an instantaneous change in population size at a single point in time *t*\* from constant size *N* to constant size *κ_N_*, where *κ* is a positive parameter. In other words, the population size over time is a step function *N_κ_* such that *N_κ_*(*t*) = *N, t* < *t*\*, and *N_κ_*(*t*) = *κN*, *t ≥ t*\*. Throughout this work we use *N* as reference size and apply an evolutionary timescale where *t* units of time correspond to [*Nt*] generations. Each individual is characterized by *L* independent sites, which corresponds to the assumption of free recombination between sites. Random mutations arrive independently and uniformly over individuals on mono-allelic sites with population mutation intensity *θ* per generation in the reference population. Hence, as long as *t* ≤ *t*\*, the mutation intensity per time unit is *θN*. Consequently, for *t* ≥ *t*\*, the mutation intensity is *kΘ* per generation and *kΘN* per time unit. Each mutation is assigned a population selection intensity *γ*.

We use the Wright-Fisher model with selection (Fisher 1930; Wright 1931) for two alleles segregating at one site to model reproduction and then study the population dynamics of the collection of all *L* independent sites. In the limit as *L* tends to infinity and *N* is large but fixed, the Poisson random field approximation (Sawyer and Hartl 1992; Kaj and Mugal 2016) applies. This means that the number of new mutations over all mono-allelic sites is approximately Poisson distributed with mean *θN_κ_* per time unit. The derived allele frequencies in a polymorphic site starting at time *s* evolve as independent Wright-Fisher diffusion processes with selection, that is, solutions of the stochastic differential equation

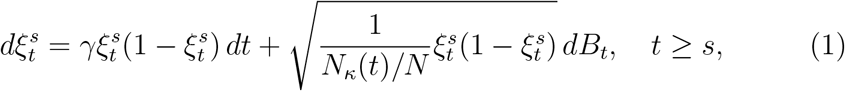

with initial value 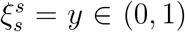, where *B_t_* is a standard Brownian motion. Here, *N_κ_*(*t*)/*N* = 1 whenever *t* ≤ *t*\* and *N_κ_*(*t*)/*N* = *κ* for *t > t*\*. We denote such a Markov process by 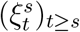 or simply (*ξ_t_*)_*t*≥0_ when the initial time is *s* = 0. Furthermore, let 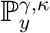 and 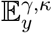 be the law and expectation of processes (*ξ_t_*)_*t*_ that start in *y*, have selective pressure *γ*, and evolve in a population of size *κN*. Let *τ*_1_ be the time to fixation of the derived allele. The fixation probability is given by (Kimura 1962)

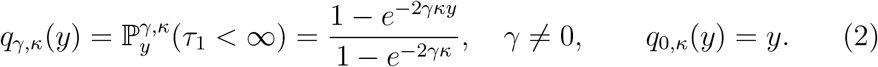

As *N* → ∞, the scaled fixation rate emerges as

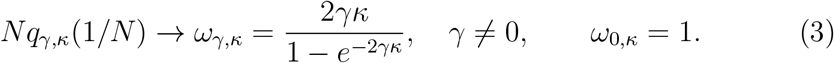

This means, in an equilibrium population of size *κN*, the fixation rate ratio of a class of selected (with selective pressure 7) and neutral mutations in the limit equals *ω_γ,κ_*.

Returning to the Poisson random field setting, the allele frequencies are represented by Poisson points (*s,ξ^s^*) on the collection of sites according to the Poisson distribution with intensity *θN_κ_*: once such a mutation event takes place at a certain time *s*, a path 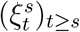 is initialized at frequency 1/*N_κ_*(*s*). We fix *t*\* > 0 and represent the state of the Poisson random field at time *t* as a random measure 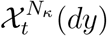 (dy) on (0,1]. For *t* ≤ 0, however, we will ignore fixations but start from polymorphic frequencies on (0,1) at *t* = 0. A visualization of the setup is presented in Fig. 1. It is known that the aggregate of all mutations from the infinite past in the ancestral population together build up a Poisson measure in steady state. Let

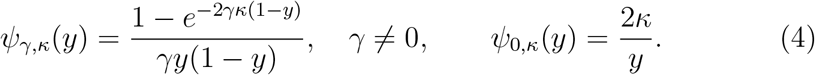

**Figure 1:**
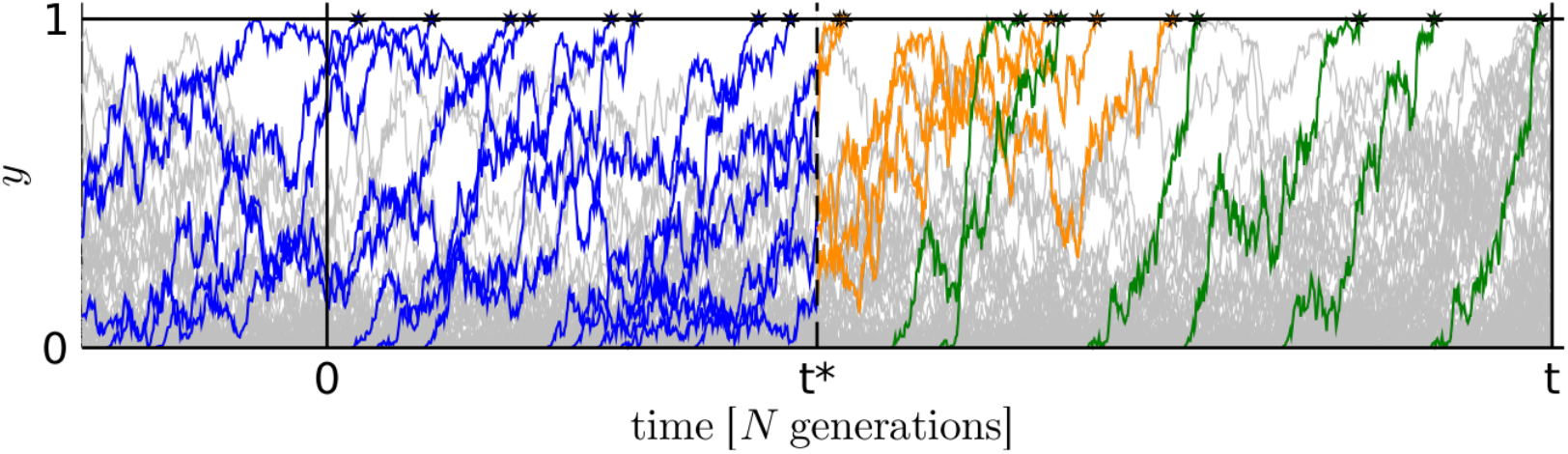
Polymorphisms and fixations in [0, *t*]. Vertical lines as for example at *t*\* represent a snapshot of the population dynamics and the distribution of allele frequencies at that specific point in time are summarized in the AFS. Polymorphisms in grey go to extinction or still segregate at *t*. Polymorphisms leading to a substitution are colored: fixations before *t*\* (blue), fixations after *t*\* emerging from segregrating polymorphisms at *t*\* (orange), and fixations of mutations that arose after the change in population size (green).

Now the relevant initial distribution at *t* = 0 for our model, that is 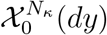, is a Poisson measure with intensity measure *ω*_*γ*,1_ *ψ*_*γ*,1_(*y*)*dy* on (0,1). The initial distribution at *t* = 0 plus the arrival of new mutations during (0,*t*] together preserve the Poisson distribution which is invariant as long as the population size does not change, i.e. for 0 < *t* ≤ *t*\*. To account for fixations during [0,*t*] we also include the singular contribution at 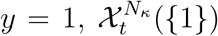, *t* ≥ 0, which is a Poisson counting process with time-inhomogeneous intensity.

Formally, we construct our model 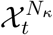 using stochastic Poisson integrals. The detailed presentation and most of the technical aspects are deferred to the appendix sections. For 0 < *t* ≤ *t*\*,

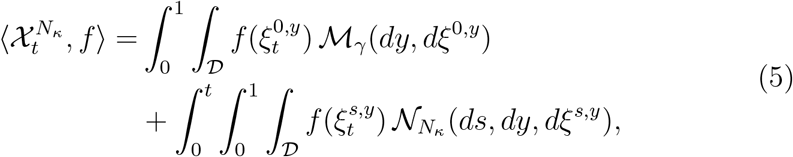

for suitable functions 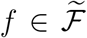, specified in Appendix A, satisfying sufficient conditions for these integrals to be well defined. In particular, *f* (0) = 0. The class 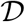 is the path space for the diffusion processes *t ↦ ξ_t_*, consisting of functions *g*: ℝ → [0,1] which are right continuous and have left limits. Moreover, 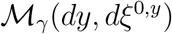 is a Poisson random measure on 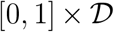 with intensity 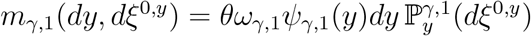 and 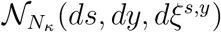 is a Poisson random measure on 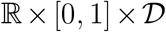 with intensity measure 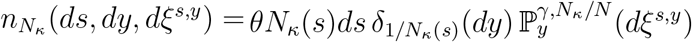. The first term in Eq. (5) represents the family of ancestral allele frequencies with initial values at *t* = 0 given by the Poisson measure 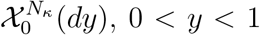. The second term contains additional allele frequencies sparked off by mutations during (0, *t*]. Similarly, the collection of paths at some time *t > t*\* consists of two contributions,

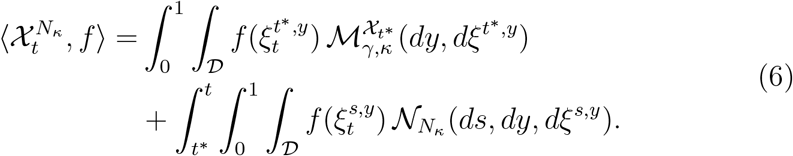

Here, conditional on 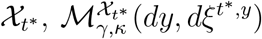 is a Poisson random measure on 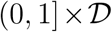 with intensity 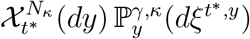. This term represents the fate of the allele frequencies extending beyond *t*\* of all alleles, polymorphic or fixed, which were present at *t*\*. The additional term again covers new mutations taking place subsequent to the change in population size.

### The nonequilibrium AFS

The AFS accounts for the collection of all derived allele frequencies across sites at a fixed point in time. The spectrum of allele frequencies *y*, 0 < *y* < 1, represents the average intensity of attained frequency values at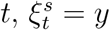 for some *s ≤ t*, compare Fig. 1. In our approach we also include alleles which have reached fixation during [0, *t*]. As a reference case we begin with the equilibrium AFS, which arises as the scaled limit of expected values for the case of a fixed size population, say *κN*. Then,

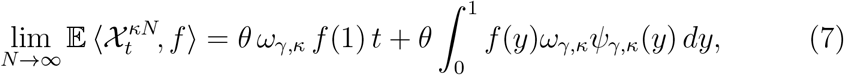

see Lemma 3 iii) in Appendix A. In particular, taking *κ* =1, the limiting expected value of relation (5) is given by this linear function, as long as *t ≤ t*\*. The linear term in t represents the effect of constant rate fixations and the integral term independent of time represents the steady-state spectrum of polymorphic frequencies.

In order to analyze the nonequilibrium AFS caused by applying population size *N_κ_*, we derive the limiting expected values of each term in Eq. (6), see Theorem 1, Appendix B. The limiting expected contribution from the ancestral component in Eq. (6), mutations which occurred prior to *t*\*, equals

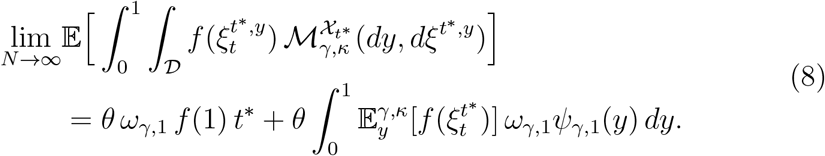

compare Eq. (B.1) in Theorem 1. Similarly, the limiting expected value of the latter contribution in Eq. (6) yields the nonstationary build-up AFS (Kaj and Mugal 2016, Theorem 1), that arises from a completely mono-allelic population. In Lemma 3 i) together with Theorem 1 we show

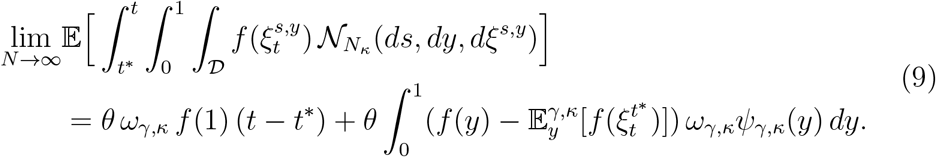

While in these representations we do not see directly a spectrum of frequencies y with explicit weights affecting *f* (*y*), we do see indirectly the timedependence effect due to the nonequilibrium framework. By adding up both contributions we obtain our main result regarding the nonequilibrium AFS, as

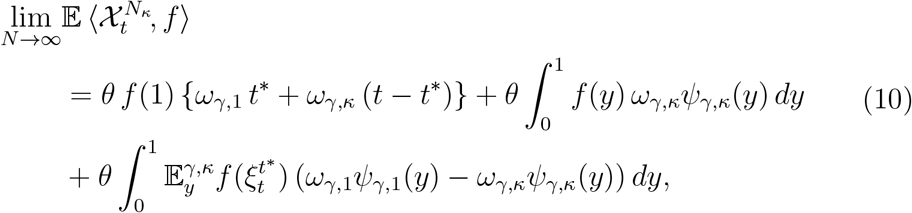

see Theorem 1 in Appendix B for a detailed derivation.

We make two remarks: for 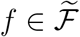 with the additional property *f* (1) = 0, the time-dependent AFS provides a bridge between the “marginal” limit spectra

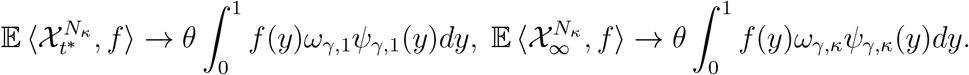

For the case of neutral evolution, *γ* = 0, we have *ω*_0,1_*ψ*_0,1_(*y*) – *ω*_0,*κ*_*ψ*_0,*κ*_(*y*) = 2(1 – *κ*)/*y* and hence, for 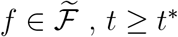, and *N* → ∞,

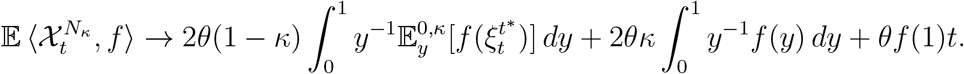

Application of selected functions f to the nonequilibrium AFS in Eq. (10) allows to retrieve expressions for relevant summary statistics, such as nucleotide diversity and the fixation rate.

### The ratio of nucleotide diversity during nonequilibrium

We derive and investigate the ratio of nucleotide diversity, *π_N_/π_S_* (Nei and Li 1979), in a population undergoing a change in population size according to *N_κ_*. Nucleotide diversity measures the number of pairwise differences, which entails integrating the specific function *f*_pw_(*y*) = 2*y*(1 – *y*) with respect to the AFS. As a reference case we observe that during equilibrium in a population controlled by a size parameter *κ* and a fixed selection coefficient *γ* ≠ 0, we have by Eq. (7) with *f*_pw_(1) = 0,

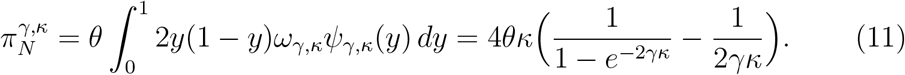

More generally, by applying Eq. (10), we obtain the time-dependent non-synonymous nucleotide diversity measure 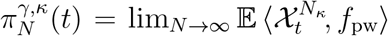 in nonequilibrium.

In order to allow for variation in selection across sites for the nonsyn- onymous diversity, we integrate the previous expressions over a distribution of fitness effects (DFE). We will denote the random variable generating the *γ*-values by 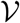 and assume it has a continuous density function 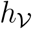. Be-cause of the presumed rarity or negligibility of advantageous mutations at the genome-wide level within the nearly neutral theory, we focus on weak and strong purifying selection following Eyre-Walker *et al.* (2006); Loewe and Charlesworth (2006); Galtier and Rousselle (2020). A common choice of DFE in this scenario is the negative gamma distribution. The density function is

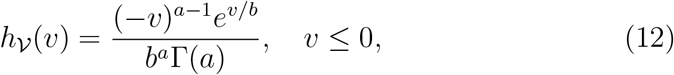

with shape parameter *a* > 0, scale parameter *b* > 0, and mean — *ab*. Integration of the expression in Eq. (11) and 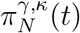 over this density yields an averaged diversity measure 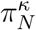. Taken together it holds

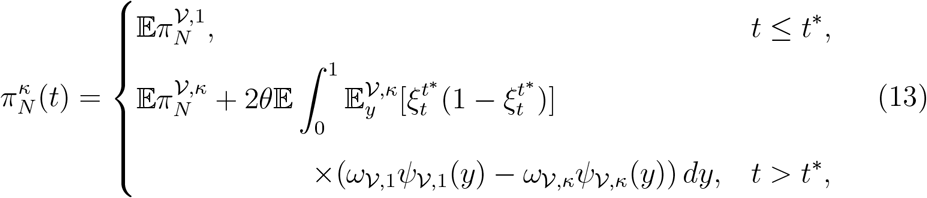

see Appendix C for details. The expectations in the above expression are used to indicate integration over the DFE. We observe that 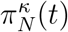 approaches a new equilibrium, 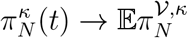 as *t* → ∞, since 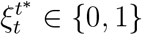 for *t* → ∞. For the case of neutral evolution, *γ* = 0, the result simplifies considerably and we obtain the synonymous diversity as

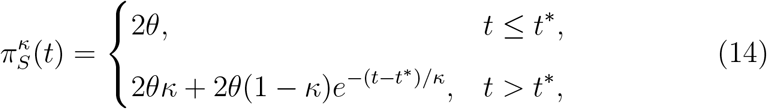

with 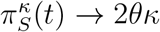 as *t* → ∞. The ratio of nonsynonymous and synonymous diversity, which we denote 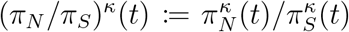, is also time dependent and determined by Eqs. (13) and (14). The ratio in equilibrium will be denoted (*π_N_/π_S_*)^κ^.

### The fixation rate ratio during nonequilibrium

The nonsynonymous to synonymous fixation rate ratio of selected and neutral mutations (Goldman and Yang 1994; Muse and Gaut 1994), *ω^κ^*(*t*), is determined by the ratio of the number of nonsynonymous and synonymous fixations in a finite interval. We therefore consider the number of fixations in the population during [0,*t*] with a change in population size as before given by *N_κ_*. To account for fixations in the random field setting, we wish to count all Poisson points (*s,ξ^s^*) such that 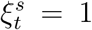. In other words, we evaluate the indicator function 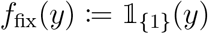 at the nonequilibrium AFS, Eq. (10). Hence, the number of fixations in the population during [0,*t*] with a change in population size given by *N_κ_* is 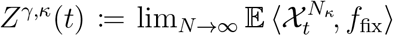. The decomposition of 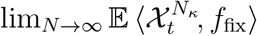 into the ancestral contribution in Eq. (8) and the buildup in Eq. (9) allows for matching the different categories of fixations in Fig. 1 with the corresponding analytic representation: the first term in Eq. (8) reflects fixations (in blue) appearing during [0,*t*\*], whereas the second part corresponds to fixations (in orange) during [*t*,t*] for which the mutation happened before *t*\*. The buildup component in Eq. (9) accounts for fixations (in green) during [*t*, t*] for which the mutation occurred after *t*\*.

To obtain an explicit representation of *Z*^*γ,κ*^(*t*), we note that *f*_fix_(1) = 1 and that the expectation operator applied to *f*_fix_(*y*) can be rewritten in terms of the fixation time distribution,

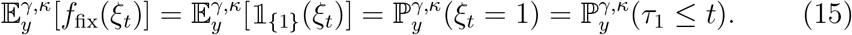

Thus,

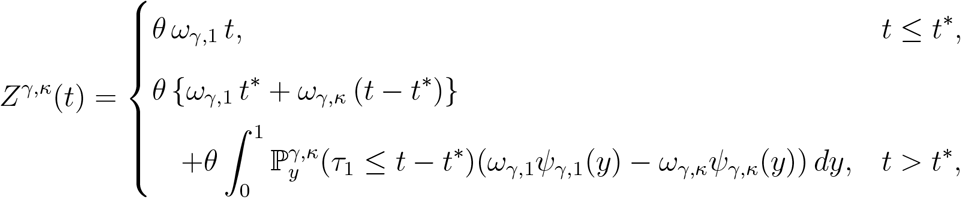

compare Appendix D for technical details. Fixations that originate from nonsynonymous mutations are averaged over the DFE in Eq. (12); for synonymous fixations *γ* is set to zero. Finally, the fixation rate ratio in nonequilibrium is defined as

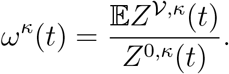

The nonequilibrium quantity *ω^κ^*(*t*) is consistent with the equilibrium fixation rate ratio *ω_γ,κ_* stated in Eq. (3), since 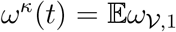 for *t ≤ t*\* and 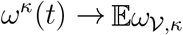 for *t* → ∞.

We note that measures modeled in this study are population functionals— in contrast to sample functionals which can be viewed as estimators of population functionals. In case of (*π_N_/π_S_*)^*κ*^(*t*) the population and sample functional are equal. When using nonsynonymous over synonymous sequence divergence as estimate for the fixation rate ratio *ω^κ^*(*t*), sample functional and population functional are not equal but converge for *t* → ∞ (Wolf *et al.* 2009; Mugal *et al.* 2014, 2020).

### Stochastic simulations

For performing stochastic simulation of paths (*ξ_t_*)_*t*_, we apply the discrete Wright-Fisher model with selection to a population of size *N*. It suffices to simulate paths for the reference population, since a polymorphism in a population of size *κN* evolves as in the reference population but with time scaled by *κ*. Paths are simulated over a maximum of *n* generations using the binomial Wright-Fisher sampling with selection. The selection coefficient in the discrete setting is obtained from the relation *s = γ/N*.

For the distribution of the time to fixation, 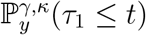, we generate *m*_*τ*1_ paths for each tuple (*y, γ*). If the derived allele does not get fixed, the time to fixation is set to infinity. Otherwise the fixation time is set to the generation it got fixed. Finally, the distribution function of the time to fixation on the evolutionary timescale is obtained by scaling generations with *N*. For the expected value 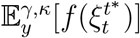 in the nonequilibrium AFS we simulate and average over m paths for each triplet (*y,γ*, (*t – t*\*)/*κ*).

Parameters that we used are *N* = 1000, *n* = 20.000, *s* ∈ [—1,0] or equivalently *γ* ∈ [–1000, 0], *m*_*τ*1_ = 100.000 (if *γ* = 0) and *m*_*τ*1_ = 10.000 (if *γ*< 0), respectively, *m*_ξ_ = 1000.

### Data availability statement

Code used to implement the stochastic simulations can be found on Git-Lab.

## RESULTS AND DISCUSSION

### Measures of natural selection in a nonequilibrium population

We study the behavior of measures of natural selection as functions of time after a change in population size. Figure 2A shows the behavior of (*π_N_/π_S_*)^*κ*^(*t*) for *κ* ∈ {0.1,0.25, 0.5,1,2, 4}. The time it takes to reach the new equilibrium depends on the direction of change in population size: the new equilibrium is reached quickly in case of a reduction (*κ* < 1), whereas for an increase in size (*κ* > 1) it takes longer. Also the extent of change determines how fast (*π_N_/π_S_*)^*κ*^(*t*) attains the new equilibrium value. The more the population size is reduced, the faster (*π_N_/π_S_*)^*κ*^(*t*) reaches its new equilibrium value; the more a population increases in size, the longer it takes. Also, given a DFE restricted to deleterious mutations, (*π_N_/π_S_*)^*κ*^(*t*) is negatively correlated with population size as predicted by the nearly neutral theory of molecular evolution.

**Figure 2:**
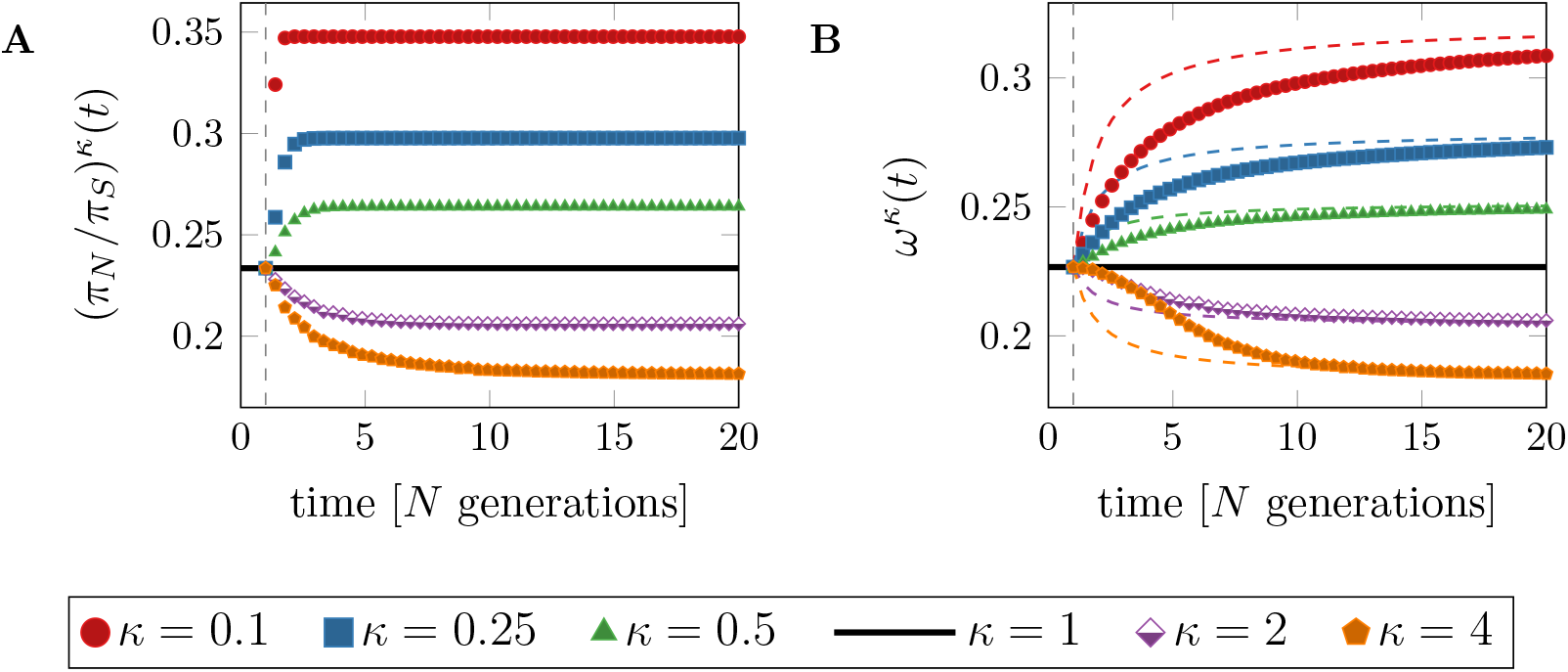
Measures of selection for different values of *κ* as functions of time. Panel A: the ratio of nonsynonymous and synonymous diversity (*π_N_/π_S_*)^*κ*^(*t*). Panel B: the fixation rate ratio *ω^κ^*(*t*). For comparison, colored, dashed curves represent the weighted fixation rate ratio 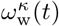. Vertical, dashed lines indicate the time *t*\* = 1. Parameters: *θ* = 1, and *a* = 0.15 and *ab* = 2500 for the DFE.

The behavior of *ω^κ^*(*t*) after a change in population size is depicted in Fig. 2B and is similar to the behavior of (*π_N_/π_S_*)^*κ*^ (*t*). The ratio decreases for *κ* > 1, which means that less deleterious nonsynonymous mutations went to fixation—in accordance with observations about selection acting more efficiently in larger populations. However, *ω^κ^*(*t*) is an accumulative measure over the time interval [0,*t*], while (*π_N_/π_S_*)^*κ*^(*t*) reflects a *snapshot* of the strength of selection at time *t*. As a consequence, *ω^κ^*(*t*) takes longer to reach its new equilibrium than (*π_N_/π_S_*)^*κ*^ (*t*).

Another model to capture the impact of a change in population size on *ω^κ^*(*t*) can be achieved by the weighted sum of the ancestral and the new equilibrium value (Eyre-Walker 2002). In our notation this reads 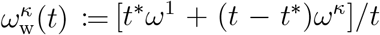 for equilibrium values *ω*^1^ and *ω^κ^*, respectively. This means, the approach based on the weighted sum does not explicitly model nonstationarity. To visualize the difference between *ω^κ^*(*t*) and 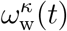, the weighted fixation rate ratios are included as dashed lines in Fig. 2B. The nonequilibrium model derived in this study shows that *ω^κ^*(*t*) takes longer to react to the change in population size and to reach the new equilibrium value compared to the approach of weighting the equilibrium values. This illustrates that ignoring the nonequilibrium phase leads to an underestimation of the time until the new equilibrium is reached.

Note that Fig. 2B shows the fixation rate ratio for a change in population size at time *t*\* = 1. Changes at other time points can change the behavior severely. A change at *t*\* = 0 for example, would lead to 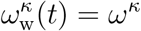 without reflecting any influence of the ancestral population. On the other hand, if the change in population size happens more recently in time, the contribution of the ancient population size becomes more pronounced (see Appendix Fig. E.6).

### Proxies of effective population size as measures of genetic drift

In order to evaluate the efficacy of selection during nonequilibrium we relate the above-introduced measures of selection to estimates of the effective population size. Since there are various ways to define *N*_eff_, it is funda-mental to discuss the differences and to assess which of the definitions are relevant to relate to *ω^κ^*(*t*) and (*π_N_/π_S_*)^*κ*^(*t*) in our modeling approach. The most commonly considered concepts of *N*_eff_ among others are variance and inbreeding (Wright 1931, 1938; Crow 1954; Crow and Kimura 1970), coalescent (Lynch and Conery 2003), and eigenvalue effective population size (Ewens 1969, 1979, 1982). The properties, that these concepts aim to model, are the variance in allele frequencies over time due to random genetic drift, the average inbreeding coefficient, the rate of coalescence of neutral alleles, and the leading non-unit eigenvalue of the allele frequency transition matrix.

We here focus on the harmonic mean effective population size over [0, *t*] (Wright 1938; Karlin 1968) and the pairwise synonymous nucleotide diversity (Lynch and Conery 2003; Wakeley and Sargsyan 2008; Ellegren and Galtier 2016). The harmonic mean effective size over [0,*t*] is a representative of variance effective population size and defined as the average of genetic drift over the time interval [0, *t*] with *t*\* ∈ [0, *t*],

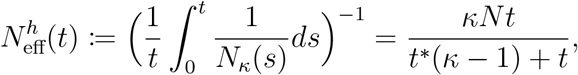

with 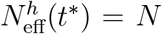, and 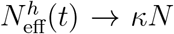 for *t* large. This means the ancestral population size *N* loses its influence on 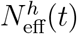 the further in the past the change took place. Overall the quantity is mainly controlled by the smaller population size (Wright 1938). If *κ* is constant over [0,*t*], then 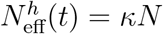. Also, in view of the genetic drift term in Eq. (1)—the variance term of the SDE—the parameter *κ* at time *t* multiplied by *N* can be seen as variance effective population size at time *t*.

The scaled pairwise synonymous diversity, 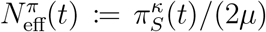, where *μ*: = *θ/N* is the mutation rate per generation and individual, is an estimate of effective population size based on nucleotide variation, often also perceived as coalescent effective population size (Lynch and Conery 2003; Wakeley and Sargsyan 2008). We note that the definition of coalescent effective population size is not consistent in literature (Sjödin *et al.* 2004). We here follow the definition of Lynch and Conery (2003); Wakeley and Sargsyan (2008).

With the two measures, 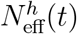 and 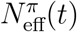, of effective population size at hand, we investigate and compare how a change in population size is reflected in each of them. We consider two scenarios, an ancient change at *t*\* = 1 and a recent change at *t*\* = 18. For this purpose, 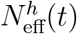 is plotted against 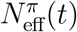 as parametric curve of time, *t ≥ t*\*, for an ancient (Fig. 3A) and recent (Fig. 3B and Appendix Fig. E.7) change in population size. The nonequilibrium behavior is compared to the equilibrium values, i.e. 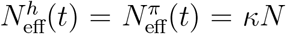, as functions of *κ*. To explain Fig. 3 more explicitly, starting from a common initial equilibrium value at time *t*\* indicated as black cross on the dashed equilibrium line, the relationship between 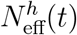 and 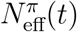 is shown for different values of *κ* for *t*\* < *t* < 20 as colored curves.

**Figure 3:**
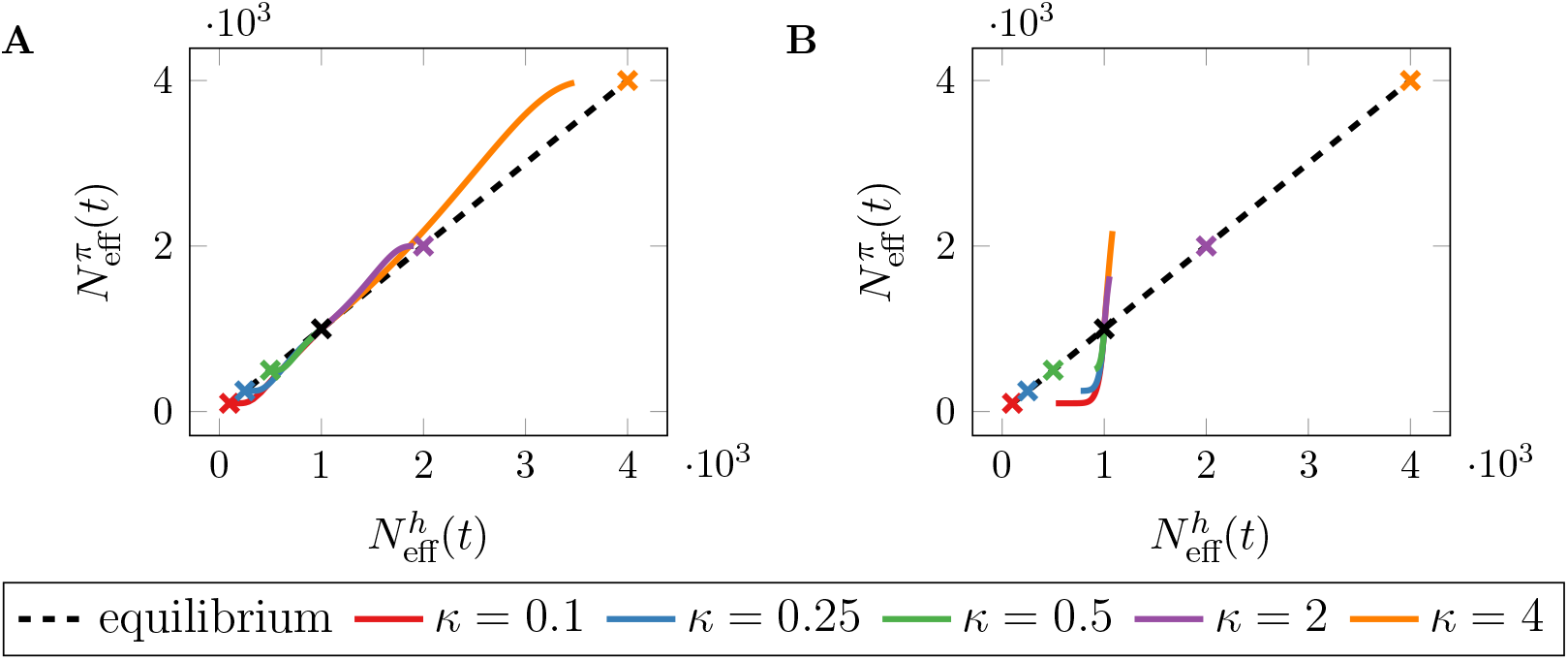
The effective population size based on nucleotide variation, 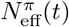, versus the harmonic mean effective population size, 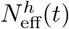, as parametric curves of time, *t*\* ≤ *t* ≤ 20, for different values of *κ*. Panel A shows an *ancient* change in population size at *t*\* = 1, panel B a *recent* change at *t*\* = 18. Black, dashed lines show the expected behavior in equilibrium populations. Black crosses at 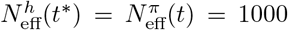 mark the time of change in population size, *t*\*. Colored crosses indicate the expected equilibrium value for each *κ*. Parameters *θ* =1 and *N* = 1000.

For an ancestral change, it seems that both proxies mirror the change in size to a large degree. However, 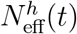 reaches the new equilibrium value more slowly compared to 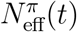. This holds in particular for *κ* > 1, leading to the presumption that the more a population increases, the slower the new equilibrium is reached and vice versa. For a recent change in population size, the difference between the two estimates is much more striking and we observe that there is hardly any correlation between them. The proxy 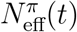 reacts quickly to a change in population size, as expected for a measure relevant at the microevolutionary timescale. The harmonic mean effective population size, 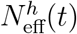, reacts more slowly and its variation is very narrow in comparison to 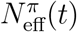. This suggests that 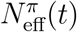 reflects a recent change in population size better than 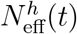. Considering the analytical definitions of the two estimates, one can additionally state that changes which happened before time zero, i.e. *t*\* ≤ 0, will not at all be captured by 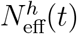 but potentially by 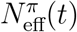 given *t* is small enough such that the AFS has not reached its new equilibrium.

### Relationship between effective population size and measures of natural selection

We investigate the selection-drift relationship after a change in population size and compare it to the equilibrium behavior. Therefore, we relate the two estimates 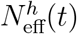 and 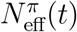 to the fixation rate ratio, *ω^κ^*(*t*), and the ratio of nucleotide diversity, (*π_N_/π_S_*)^*κ*^(*t*), as parametric curves of time after an ancient (*t*\* = 1, Fig. 4) and recent (*t*\* = 18, Fig. 5) change in size. To evaluate the discrepancy between the nonequilibrium behavior and the expected equilibrium balance between genetic drift and selection, we indicate *ω^κ^* and (*π_N_/π_S_*)^*κ*^ as functions of *κ*.

**Figure 4:**
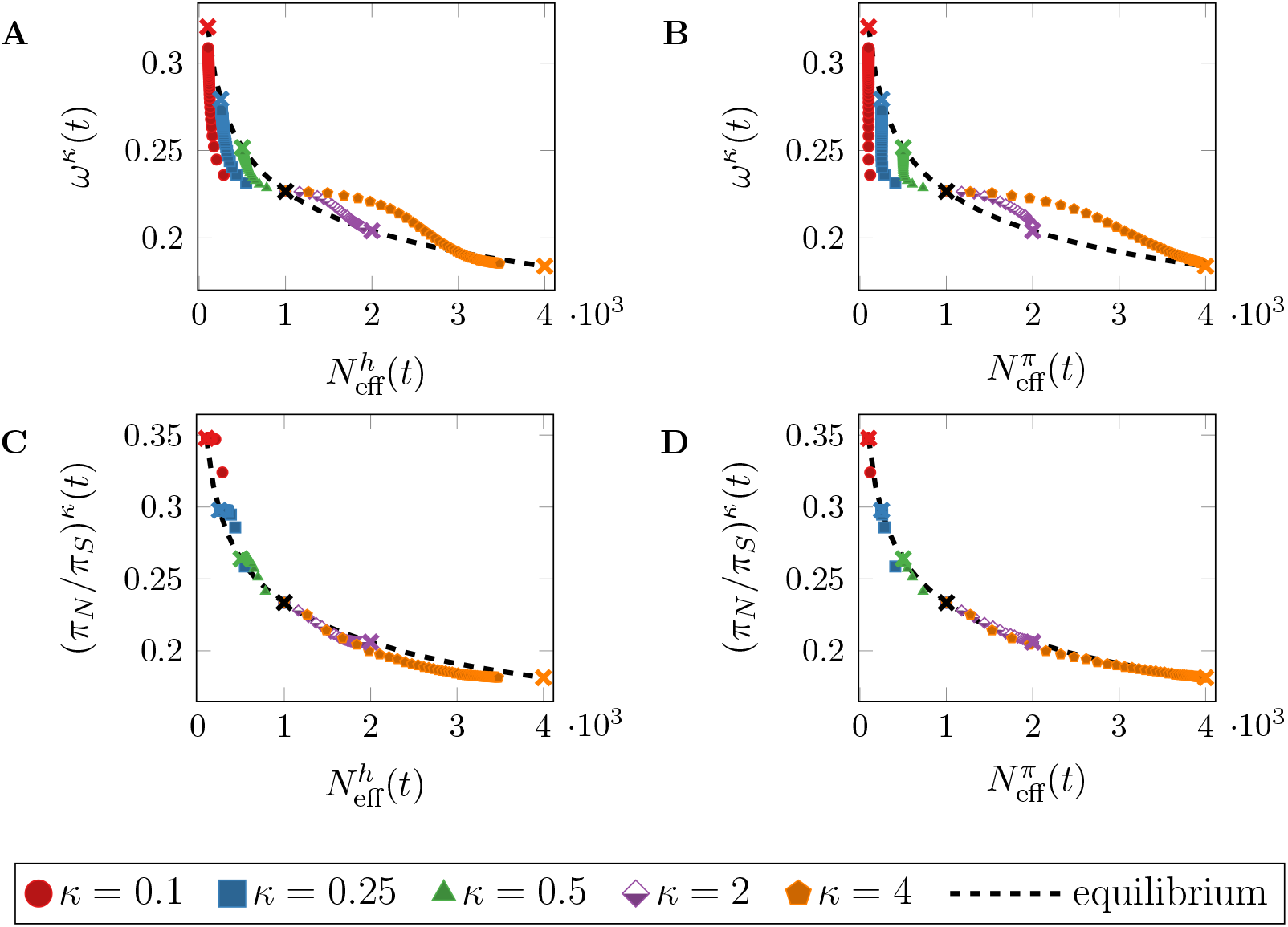
Measures of selection versus proxies of *N*_eff_ as parametric curves of time, 1 ≤ *t* ≤ 20, after an *ancient* change in population size at t* = 1 for different values of *κ*. Panels A and C: genetic drift estimated by the harmonic mean effective population size, 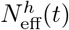. Panels B and D: genetic drift estimated by 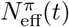. Black, dashed lines show the expected relation in equilibrium populations. Black crosses at 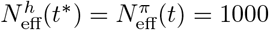 indicate the time of change in population size, *t*\*. Colored crosses indicate the expected equilibrium value for each *κ*. Parameters *a* = 0.15 and *ab* = 2500 in the DFE, *N* = 1000, and *θ* =1.

First, we consider an ancient change in population size (Fig. 4). This reveals a clear deviation from the equilibrium prediction of the nearly neutral theory for a relevant time period after the change in population size. For an increase in population size, *ω^κ^*(*t*) is larger than expected in equilibrium, implying that more nonsynonymous and hence deleterious mutations get fixed during the nonequilibrium phase than during equilibrium, see Figs. 4A and 4B. This implies that the efficacy of selection is limited during the nonequilibrium period. The reverse is true for a decrease in size. Moreover, the discrepancy between nonequilibrium and equilibrium conditions is larger, the stronger the change in population size. In addition, the deviation from the selection-drift relationship appears more pronounced when comparing *ω^κ^*(*t*) and 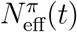 than *ω^κ^*(*t*) and 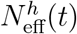. For (*π_N_/π_S_*)^*κ*^(*t*) the deviation from equilibrium is less striking—even when relating to 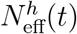 in Fig. 4C. But still, there exists a deviation which notably is the other way around than for *ω^κ^*(*t*). While for *κ* < 1 the measure *ω^κ^*(*t*) falls below the equilibrium expectation, the measure (*π_N_/π_S_*)^*κ*^(*t*) for a given *N*_eff_ is larger than in equilibrium. The reverse holds for *κ* > 1.

For a recent change in population size in Fig. 5, when using the fixation rate ratio *ω_κ_*(*t*) as measure of selection, the selection-drift relationship becomes very weak (see Figs. 5A and 5B). The lack of a clear relationship between *ω_κ_*(*t*) and *N*_eff_ can be explained by the slow reaction of *ω^κ^*(*t*) on the change in population size. Since also 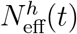 reacts slow on the change in population size, 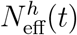 and *ω^κ^*(*t*) are both close to the ancestral value, and the lack of the selection-drift relationship is less apparent. However, it becomes clearly visible when using 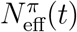 as estimate of genetic drift.

**Figure 5:**
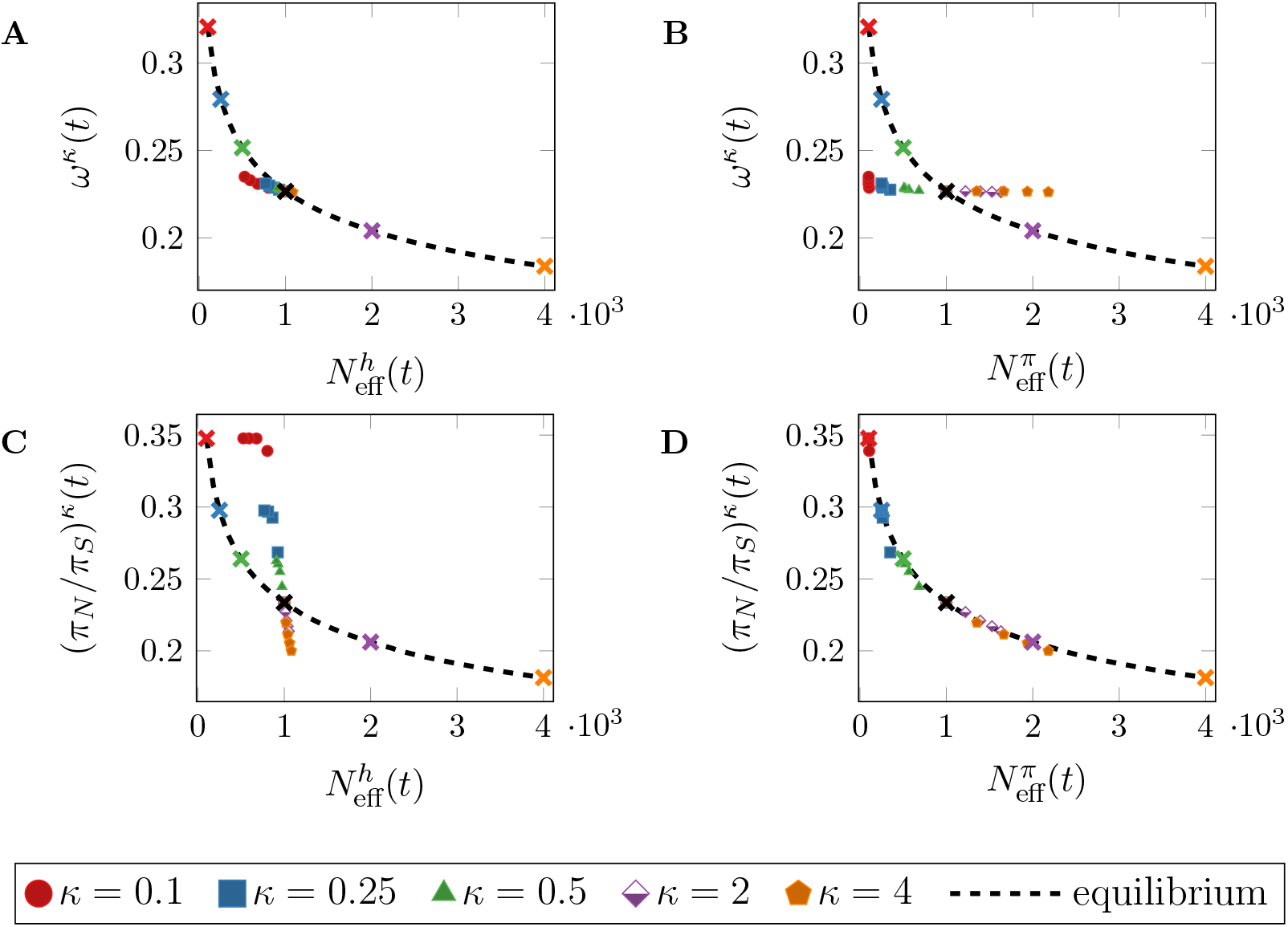
Measures of selection versus proxies of *N*_eff_ as parametric curves of time, 18 ≤ *t* ≤ 20, after a *recent* change in population size at *t*\* = 18 for different values of *κ*. Panels A and C: genetic drift estimated by the harmonic mean effective population size, 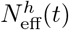. Panels B and D: genetic drift estimated by 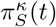. Black, dashed lines show the expected relation in equilibrium populations. Black crosses at 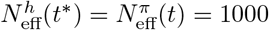 indicate the time of change in population size, *t*\*. Colored crosses indicate the expected equilibrium value for each *κ*. Parameters *a* = 0.15 and *ab* = 2500 in the DFE, *N* = 1000, and *θ* =1.

The measures (*π_N_/π_S_*)^*κ*^(*t*) and 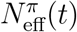 react much faster on a change in population size, which is especially eye catching in the comparison of Figs. 5A and 5D. Figure 5D shows that there is hardly any deviation from the expected equilibrium relationship since (*π_N_/π_S_*)^*κ*^(*t*) and 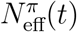 both react quickly to a change in population size and are hence close to their equilibrium expectation. This means comparison of microevolutionary measures of natural selection and genetic drift recover the equilibrium selection-drift relationship quickly. In contrast, comparison of macroevolutionary measures of natural selection and genetic drift (Fig. 5A) appear not applicable to capture recent changes in population size. If a microevolutionary measure is compared with a macroevolutionary measure after a recent change in population size, this can essentially lead to a lack of a selection-drift relationship (Fig. 5B), or stronger observed variation in natural selection than observed variation in genetic drift (Fig. 5C), dependent on the choice of combination.

Overall, we can recognize that the prediction of the nearly neutral theory, i.e. a negative correlation between measures of selection and population size, holds even after a change in population size. But the strength of relationship is clearly influenced dependent on what measures of selection and *N*_eff_ are chosen. The analysis and discussion of the various approaches to *N*_eff_ confirm that it is not trivial to choose an appropriate estimate that reflects the population behavior. Our results suggest that 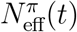 and (*π_N_/π_S_*)^*κ*^(*t*) as microevolutionary measures reflect a recent change in population size well but quickly loose the knowledge about it. In contrast, the macroevolutionary measures 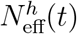 and *ω^κ^*(*t*) need more time to react to the change but are longer influenced by the ancestral population size. We can further conclude that, in order to investigate the selection-drift relationship, it seems advisable to correlate microevolutionary measures of *N*_eff_ with microevolutionary measures of selection in case of a recent change in population size. For an ancient change, microevolutionary measures of *N*_eff_ should be correlated with microevolutionary measures of selection and macroevolutionary measures of *N*_eff_ with macroevolutionary measures of selection. Moreover, it could be of interest to investigate and compare the selection-drift relationship of micro- and macroevolutionary measures. If the observed relationships show a dif-ferent behavior, this could be indicative of a nonequilibrium condition.

Obviously, data availability is an important prerequisite for such an analysis. Microevolutionary measures of genetic drift and natural selection can be directly computed based on population re-sequencing data *(Muyle et al.* 2020). On the other hand, in practice 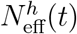 as a macroevolutionary measure of *N*_eff_ is difficult to assess based on genomic data. Instead, life-history traits, such as body mass or propagule size, are commonly used as proxies for long-term *N*_eff_ (e.g. Waples 2016; Tabak *et al.* 2018; Kutschera *et al.* 2020). The lack of a distantly related reference species can further hinder the computation of macroevolutionary measures of natural selection (Mugal *et al.* 2020; Muyle *et al.* 2020). Such limitations are often unavoidable, but our results highlight that it is important to be aware of the measures used in different studies when making comparisons among studies. When comparing the relationship between the efficacy of selection and the effects of genetic drift among different studies, as for example asexually and sexually reproducing species, or domesticated versus wild species, it could therefore be advisable to take the nature of measures of effective population size and natural selection into account (Brandvain and Wright 2016).

## Conclusion

Addressing our initial question of how a change in population size affects (*π_N_/π_S_*)^*κ*^(*t*) and *ω^s^*(*t*), both measures show a similar behavior but differ in how quickly they react to the change. For a population decrease, both ratios increase and vice versa, but (*π_N_/π_S_*)^*κ*^(*t*) as a microevolutionary measure reacts much faster to the change than the macroevolutionary measure *ω^κ^*(*t*). Second, we asked if the selection-drift relationship predicted by the nearly neutral theory also holds in a nonequilibrium. For this purpose, we compared the relationship between measures of genetic drift and selection after the change in population size with the expected equilibrium balance. Our analytical results reveal that the expected negative relationship between population size and measures of selection, such as (*π_N_/π_S_*)^*κ*^(*t*) and *ω^κ^*(*t*), is still valid, but we observe a clear deviation from the expected equilibrium balance during nonequilibrium periods. Moreover, we conclude that since measures of micro- and macroevolution show different behaviors, it is important to be aware of this fact when making inference and comparing results among studies—different observations could arise from using measures at different evolutionary timescales.

We built our model of a nonequilibrium scenario on several simplifying assumptions of which one is the focus on a single change in population size. The advantage of such narrow focus is that interpretations can be done more easily and straightforward. On the other hand, the framework we presented here is only directly comparable to a limited number of empirical scenarios, such as (but not limited to) reductions in effective population size due to isolation of island and mainland populations (Woolfit and Bromham 2005; Wang *et al.* 2014; Leroy *et al.* 2020). In addition, it might also be relevant to study periodic changes in population size as for example done by Rousselle *et al.* (2018) using simulations. Our analytical modeling approach, especially the derivation of a nonequilibrium AFS in the Poisson random field framework, may be extended to such situations. Periodic changes in population size would entail a sequence 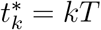, for *k* = 0,1,..., of time points of period *T* and a sequence *κ*_0_, *κ*_1_,... of size parameters making up a new step function *N_κ_*. For 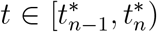, the more general nonequilibrium AFS extending Eq. (10) is

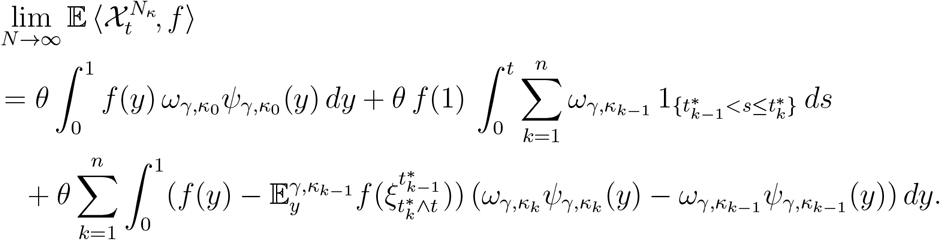

In fact, this AFS is not restricted to periodically changing environments, and hence may be used to develop the further case of allowing a prescribed, continuously varying, deterministic, scaled population size, such that *N*(*t*)/*N* → *κ*(*t*).

Apart from change in population size, there are other mechanisms that can cause a demographic nonequilibrium and affect the selection-drift balance. Examples of such mechanisms are population structure or migration. That the latter scenario can impact inference from genomic data is exemplified by the inference of estimates of gene flow in nonequilibrium conditions (e.g. Austin *et al.* 2004; Pinho *et al.* 2008). It could therefore be interesting to extend the description of the nonequilibrium AFS in order to incorporate migration and investigate its effect on the selection-drift relationship. Another evolutionary force that could impact the selection-drift relationship is recombination (Hill and Robertson 1966; Hollister *et al.* 2014; Campos *et al.* 2014). Our analytical model relies on the assumption of free recombination between sites, i.e. no linkage between sites. We show in Mugal *et al.* (2020) that measures of natural selection are affected by recombination, but that evolutionary trends of measures of selection are robust to the assumption of free recombination at least in equilibrium. However, evolutionary changes in the recombination landscape cause a deviation from equilibrium and could be a relevant topic for the future.

Finally, we restricted our analysis to purifying selection based on the argument that positive selection can be neglected due to the rarity of advantageous mutants on a genome-wide scale. This argument is still under debate (Kern and Hahn 2018; Jensen *et al.* 2018), and several empirical studies argue for a more significant role of positive selection at the genome-wide scale, (e.g. Fay *et al.* 2002; Begun *et al.* 2007; Lefebure and Stanhope 2009; Bazykin and Kondrashov 2012; Williamson *et al.* 2014). Looking at the role of positive selection could in particular be of interest when investigating the behavior of specific genes that are candidates for episodic positive selection (Schmid and Tautz 1997; Zheng *et al.* 2004; Nevado *et al.* 2019), for example immune genes (Enard *et al.* 2016) and genes involved in reproduction (Singh and Kulathinal 2000; Swanson and Vacquier 2002), rather than looking at the genome-wide average. The flexibility of our model allows for an extension to model advantageous mutations by considering for example a mixed DFE (Mugal *et al.* 2020). However, change in population size might push the population closer or further away from its fitness optimum dependent on the direction of the change, which would induce a change in the DFE. Such model extension would require a different setting, such as a mutation-selection-drift framework (Halpern and Bruno 1998; Spielman and Wilke 2015; Jones *et al.* 2016), which is a non-trivial extension.

To conclude, some of the model assumptions constitute simplifying as-sumptions that lead to a narrow focus. The crucial advantage of such a narrow focus is that exact analytical solutions can be derived and that valuable conceptual understanding can be gained based on straightforward interpretations. The framework we presented here is also flexible and allows for various modifications, which gives the opportunity to investigate different extensions in some future work.

## ACKNOWLEDGMENTS

The authors thank Martin Lascoux for valuable discussions and David Widmann for generous advice on implementing simulations in the Julia pro-gramming language. C.F.M. is funded by grants to Hans Ellegren from the Swedish Research Council (2013/08271) and Knut and Alice Wallenberg Foundation.

## A. THE AFS FOR THE EQUILIBRIUM CASE

We begin with the reference case *κ* = 1 of fixed population size and consider the Wright-Fisher diffusion 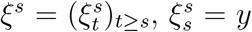, with law 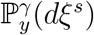. To emphasize the initial value we may write 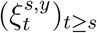. The associated expectation operator 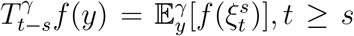, satisfies the semigroup property 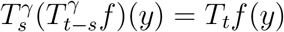, and the diffusion infinitesimal generator is the differential operator

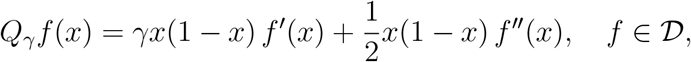

for a suitable domain 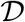 of twice differentiable functions on the unit interval. We denote by 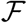 the class of real-valued bounded functions on [0,1] with *f* (0) = 0, satisfying ∫*y*^-1^|*f* (*y*)| *dy* < ∞, and put

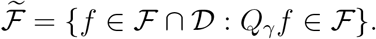

Similarly, 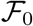 and 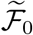 denote the further restricted classes of functions which, in addition, satisfy *f* (1) = 0. It is convenient to derive the following property first for the restricted class of functions vanishing at both boundary points, case i), and then make the necessary observations for handling the larger class of functions, as case ii).

### Lemma 1.

*For each t* ≥ 0,

i. *if* 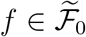 *then* 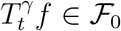; *and*
ii. *if* 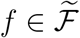 then 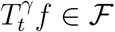.

*Proof.* For statement i) let 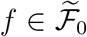. Clearly, 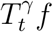 is bounded with 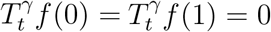. We need to show 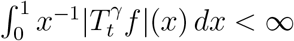.

As a consequence of Itô’s formula we have the basic relation

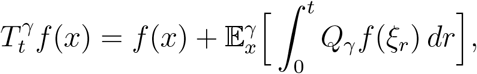

and so

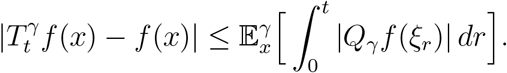

Let *τ* be the absorption time of (*ξ_t_*). Then, since 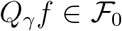 by assumption,

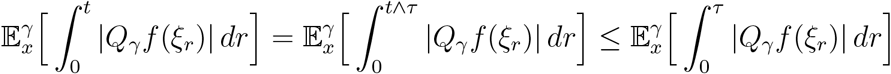

so that

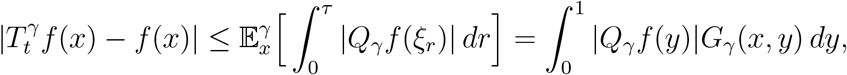

where *G_γ_*(*x,y*) is the Green function of the Wright-Fisher diffusion. Hence,

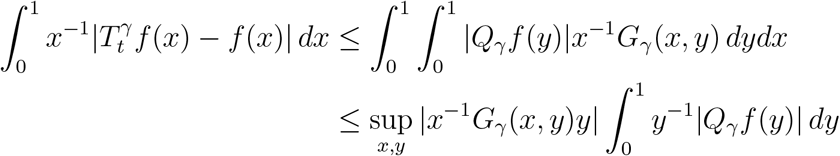

and therefore, uniformly in *t*,

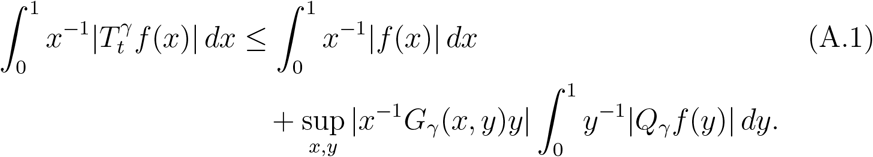

As the two integrals on the right hand side are finite by assumption, it remains to show that

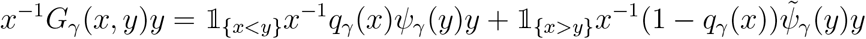

is bounded on the unit square 0 ≤ *x, y* ≤ 1. Here, *q_γ_*(*x*) is defined in Eq. (2), *ψ_γ_*(*y*) in Eq. (4), and

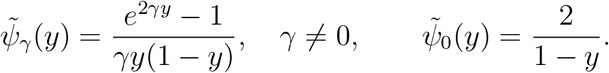

For *γ* = 0,

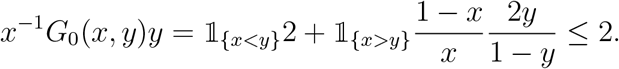

For *γ* = 0, the nonnegative function *x*^-1^ *G_γ_*(*x, y*)*y* has the representation and upper bound

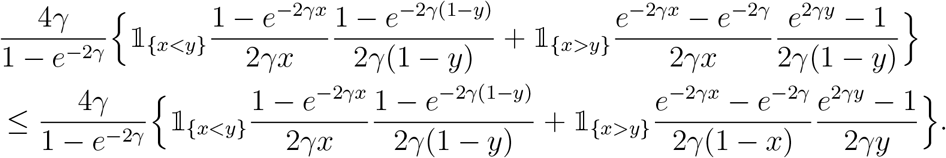

From here using straightforward estimates one obtains, for example,

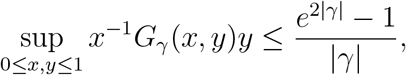

which concludes the proof that 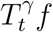 belongs to 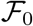.

Turning to statement ii), by using Kaj and Mugal (2016, Lemma 1, Eq. (30)), we now have

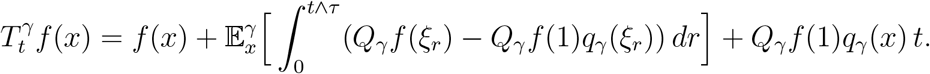

Since *q_γ_*(1) = 1, the integrand function *y ↦ Q*_*γ*_*f* (*y*) – *Q_γ_f* (1)*q_γ_*(*y*) belongs to 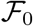. Hence we can proceed as above and obtain as replacement of Eq. (A.1),

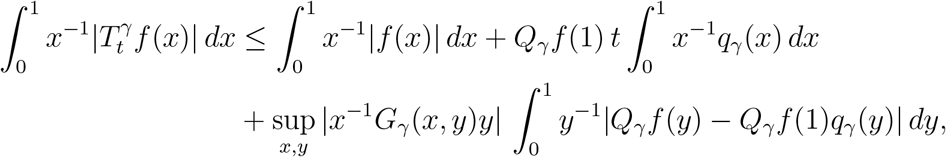

where the supremum is the same factor already treated under i). The observation

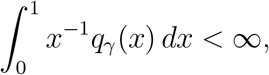

completes the proof.

The stationary AFS arises in the Poisson random field approach as a limiting intensity measure of a random Poisson measure which adds up the allele contributions from all mutations in the infinite past. Formally, let 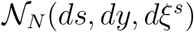 be a Poisson measure on 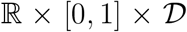 with intensity 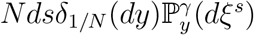, and let 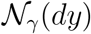 be a Poisson random measure on [0,1] with intensity measure *ω_γ_ψ_γ_*(*y*)*dy*. Note that for simplicity we assume *θ* = 1 throughout Appendix A and Appendix B. For 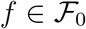, we have for fixed *t* as *N* → ∞ the convergence in distribution

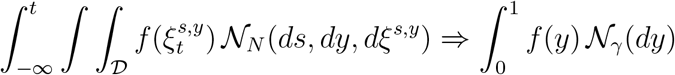

and the associated convergence of expected values

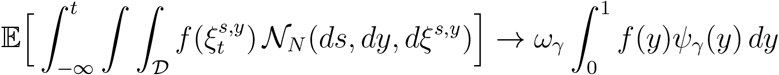

(Kaj and Mugal 2016, Section 2.4.). Each Poisson point in 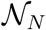 represents the allele frequency of a mutation starting at time s at a fraction 1/*N* of the population. Taking all mutations with *s ≤ t* together and considering the resulting spectrum of polymorphic sites at *t*, we obtain 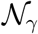.

Next, we decompose the stationary Poisson measure 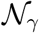 at a fixed time *t* ≥ 0, in two independent contributions by splitting up the mutation times, separating *s* ≤ 0 and 0 < *s* ≤ *t*, so that

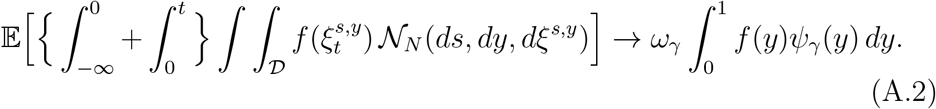

The limit of the first term in the decomposition is obtained in the following lemma, relation ii). The limit of the second term follows immediately, relation iii). Let 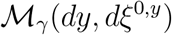 be a Poisson random measure on 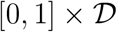 with intensity measure 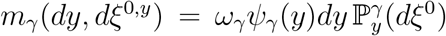. The Poisson points (*y, ξ*^0,*y*^) are paths with initial state *ξ*_0_ = *y* according to the stationary intensity *ω_γ_ψ_γ_*,(*y*)*dy* and evolve as Wright-Fisher diffusions 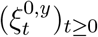.

### Lemma 2.

*For* 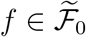 *we have as N* → ∞ *the convergence in distribution*,

i. 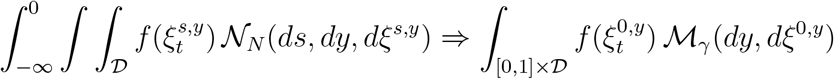 *and convergence of expected values*,
ii. 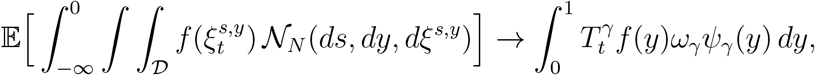 and
iii. 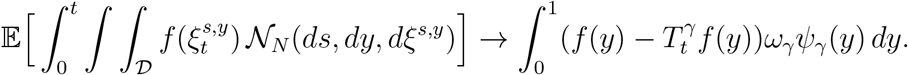

*Proof.* The logarithmic moment generating functional of the *N*-dependent Poisson integral equals

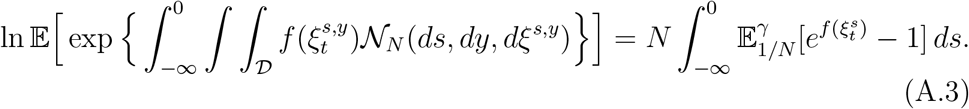

Here, conditioning on 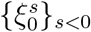,

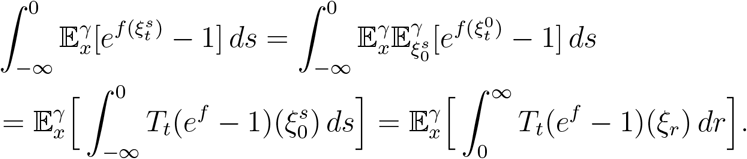

Since 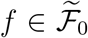 we have 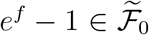. By Lemma 1 i), 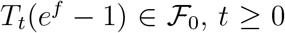.

Thus, the rightmost expression in Eq. (A.3) equals

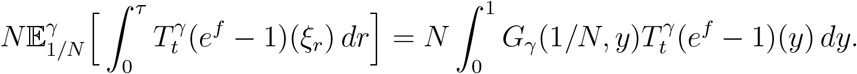

Therefore, as *N* → ∞,

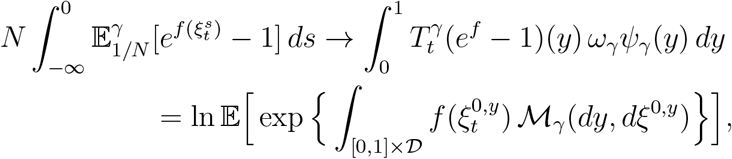

which shows convergence of the marginal distributions, for a fixed *t*. The convergence of the finite-dimensional distributions can be established as in Kaj and Mugal (2016, Proposition 2). Statement ii), which is

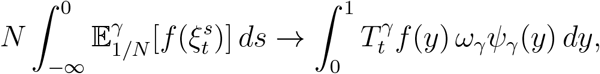

follows as above, now using that 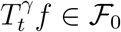. Then we obtain iii) by subtracting ii) from Eq. (A.2).

In view of Lemma 2 i), from now on we consider the allele frequency model given by the Poisson integral 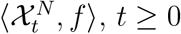, defined by

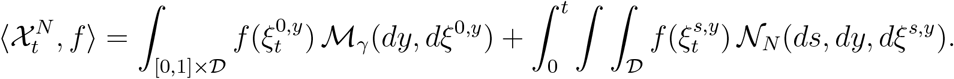

Here, the site frequency paths enter the system either at *t* = 0 according to a Poisson measure with intensity *ω_γ_ψ_γ_*(*y*) *dy*, 0 ≤ *y* ≤ 1, or at constant rate *N ds, s* ≥ 0, with initial frequency 1/*N*. In particular,

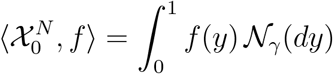

is independent of *N* and we write 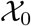. This setting allows us to analyze the effect of allele fixations, by dropping the restriction *f* (1) = 0 required for 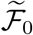. The next result shows that fixations build up at an asymptotically linear rate over time with the deviation from linearity controlled by

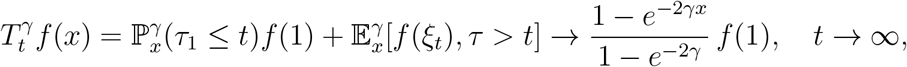

where *τ*_1_ is fixation time and *τ* absorption time.

### Lemma 3.

*For each* 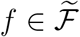 *as N* → ∞

i. 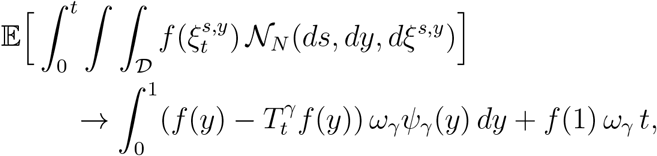
ii. 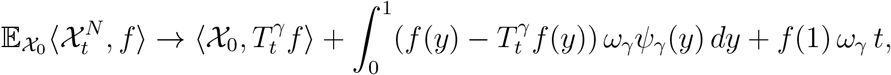
iii. 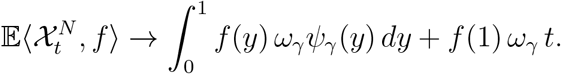

*Proof.* We begin with the basic representation of the expected value given by

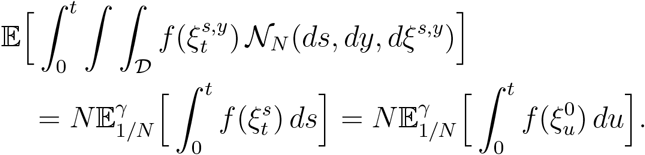

and apply the rewriting

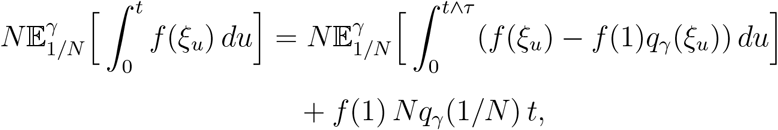

see Kaj and Mugal (2016, Section 6.3, Eq. (30)). Here, *N_qγ_*(1/*N*) → *ω_γ_*, *N* → ∞. Moreover, the function *g* defined by *g*(*y*) = *f* (*y*) – *f*(1)*q_γ_*(*y*) belongs to 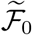 and hence, by iii) in Lemma 2,

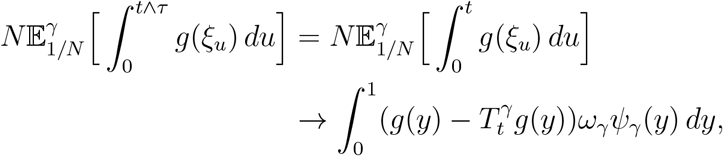

where

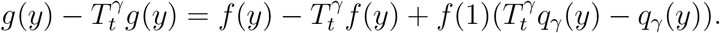

It is a well-known property of the Wright-Fisher diffusion, derived using Ito’s formula, that *q_γ_*(*ξ_t_*) is a 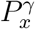-martingale, in particular 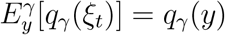. Thus 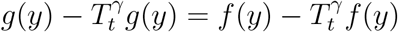, and so

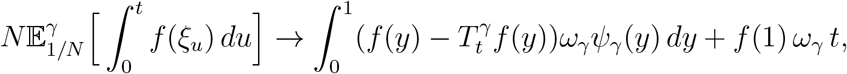

as required to show i). Statements ii) and iii) follow by observing

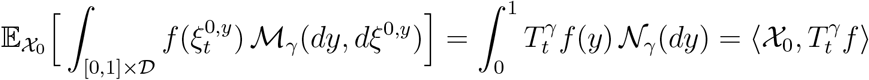

and

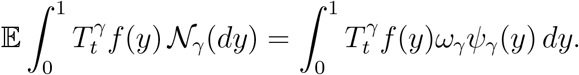

## B. THE AFS DURING NONEQUILIBRIUM AFTER A CHANGE IN POPULATION SIZE

The results in Appendix A extend to the (*γ, κ*) Wright-Fisher model with infinitesimal generator

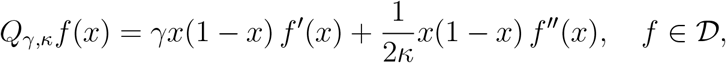

by invoking the associated expectation operator 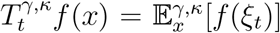, the scaled fixation probability *ω_γ,κ_*, and the intensity function *ψ_γ,κ_*(*y*), introduced in Eqs. (3) and (4). We are then in position to derive the AFS during nonequilibrium enforced by population size *N_κ_*, that is, we run the system with *κ* = 1 up to a time point *t*\* ≥ 0 and apply *κ* ≠1 from thereon. The modified Poisson random measure, denoted 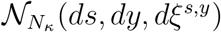, applies Poisson points for which the dynamics of the paths 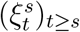 change with the current size of the population.

### Theorem 1.

*For* 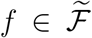 *the nonequilibrium AFS at time t ≥ t* after a change in population size at t* is given by*

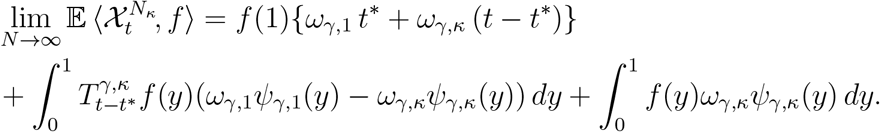

*Proof*. The allele frequencies originating from mutations before *t*\* form a stationary AFS with Poisson intensity *ω*_*γ*,1_*ψ*_*γ*,1_(*y*) dy at time *t*\*, compare Lemma 2 i) for *t* = 0. Hence,

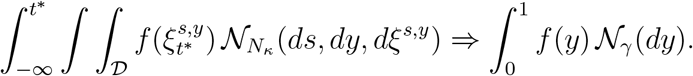

Conditioning on 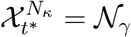, by Lemma 3 ii) (with *t*\* replacing *t* = 0 as initial time),

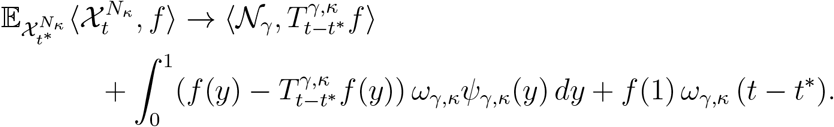

By Lemma 3 iii),

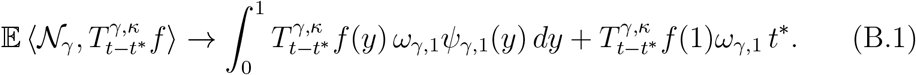

Since 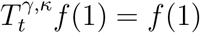 for each *t*, it follows that

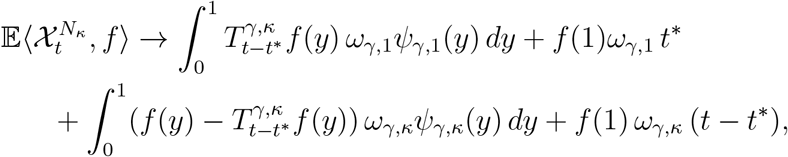

which is the desired relation and hence completes the proof.

## C. NUCLEOTIDE DIVERSITY

For computing nucleotide diversity, we apply the AFS to the function *f*_pw_(*y*) = 2*y*(1 – *y*) and note that *f*_pw_(1) = 0. For the neutral case it holds in equilibrium according to Eq. (7)

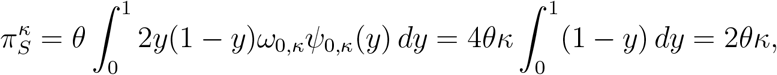

since *ω*_0,*κ*_ = 1 and *ψ*_0,*κ*_ = 2*κ/y*. The synonymous diversity during nonequilibrium is obtained by applying the AFS in Eq. (10) to *f*_pw_(*y*),

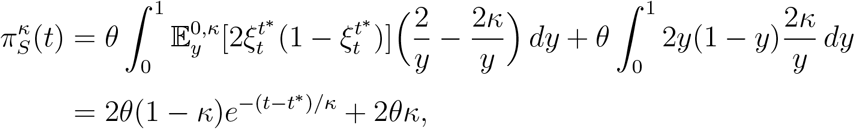

where we used Itô’s formula to compute 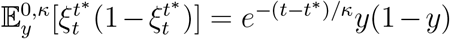. It holds 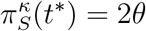 and 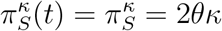 in the limit *t* → ∞.

For nonsynonymous nucleotide differences we proceed similarly but use the DFE in Eq. (12) to allow for variation in selection. The stationary case follows by integration of Eq. (11) over the DFE,

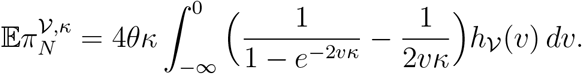

This integral is well-defined, which can be seen by looking at the Taylor expansion of the integrand around zero and noting that the moments of the gamma distribution are finite. For the behavior of *π_N_* in nonequilibrium we apply the nonstationary AFS stated in Eq. (10) to *f*_pw_(*y*), i.e.

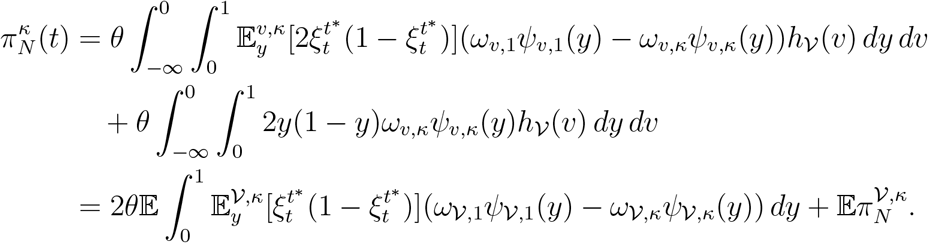

Since 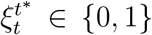 for *t* → ∞, it holds 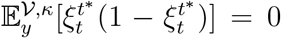 in the limit *t* → ∞, implying 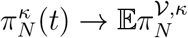. We can further explicitly write

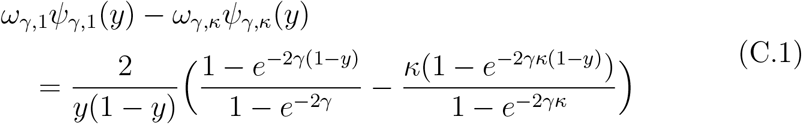

for *γ* ≠ 0, and *ψ*_0,1_(*y*) – *ψ*_0,*κ*_(*y*) = 2/*y* – 2*κ/y* = 2(1 – *κ*)/*y*.

## D. THE NUMBER OF FIXATIONS

The number of fixations before the change in population size, i.e. *Z*^*γ,κ*^(*t*) for *t ≤ t*\*, is obtained by applying the equilibrium AFS in Eq. (7) to *f*_fix_(*y*),

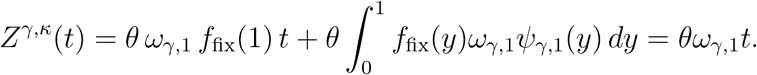

The integral term vanishes and *κ* = 1, since for *t ≤ t*\* the population size is *N_*κ*_* = *N*.

For *t > t*\*, we apply the nonequilibrium AFS in Eq. (10),

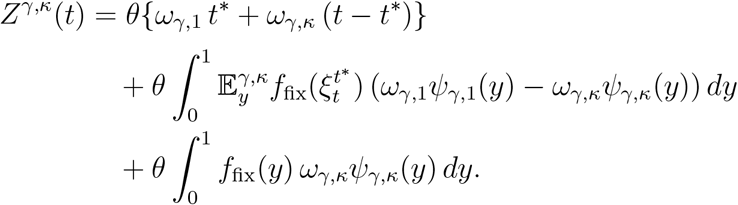

Since the last term vanishes and 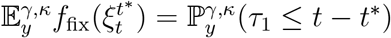, as we have shown in Eq. (15), it follows

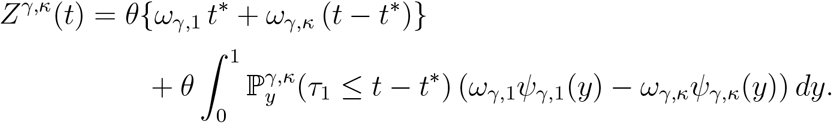

The explicit representation for *ω*_*γ*,1_*ψ*_*γ*,1_(*y*) – *ω*_*γ,κ*_*ψ*_*γ,κ*_(*y*) is given in Eq. (C.1).

## E. SUPPLEMENTARY FIGURES

**Figure E.6:**
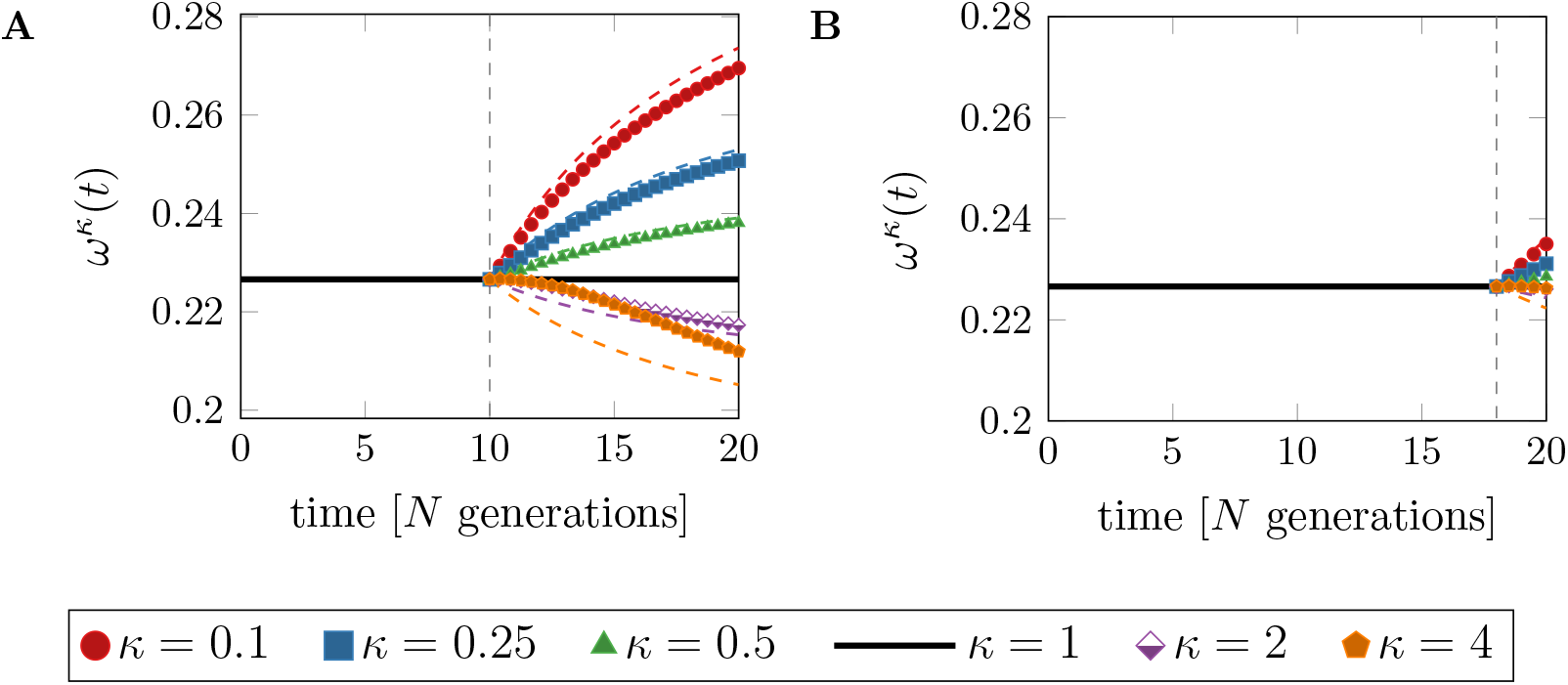
The fixation rate ratio *ω^κ^*(*t*) for different values of *κ* as functions of time. Panel A: change in population size at time *t*\* = 10. Panel B: change in population size at time *t*\* = 18. For comparison, colored, dashed curves represent the weighted fixation rate ratio 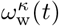. Vertical, dashed lines indicate the time *t*\*. Parameters: *θ* =1, and *a* = 0.15 and *ab* = 2500 for the DFE.

**Figure E.7:**
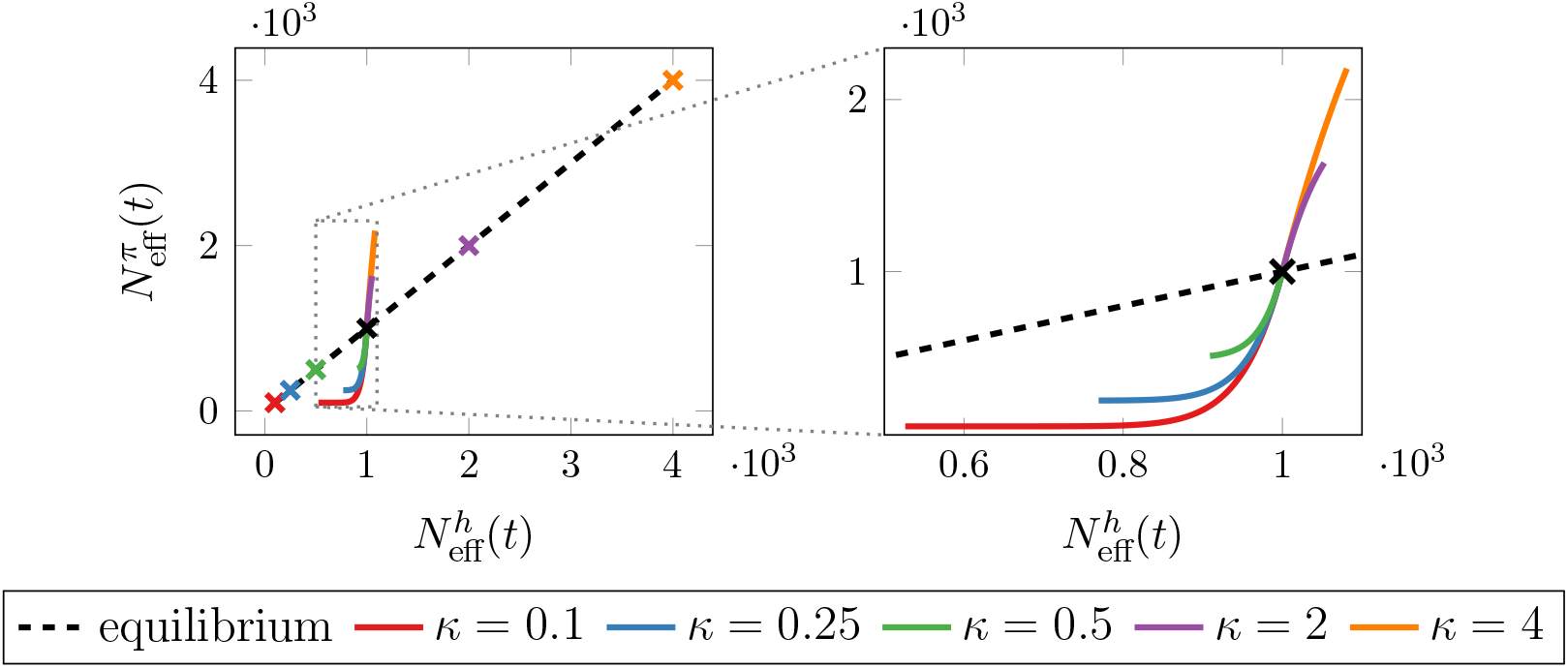
The effective population size based on nucleotide variation, 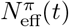, versus the harmonic mean effective population size, 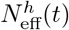, as parametric curves of time after a *recent* change in population size, *t*\* = 18 ≤ *t* ≤ 20, for different values of *κ*. Black, dashed lines show the expected behavior in equilibrium populations. Black crosses at 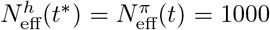 mark the time of change in population size, *t*\*. Colored crosses indicate the expected equilibrium value for each *κ*. Parameters *θ* =1 and *N* = 1000.

